# Structural extension of the human exocyst is enabled by a minimal interface

**DOI:** 10.1101/2024.10.07.617058

**Authors:** Haonan D. Xu, Mihaly Badonyi, Watanyoo Sopipong, Stuart A. MacGowan, Joseph A. Marsh, David H. Murray

## Abstract

In multicellular organisms, the machinery responsible for polarized trafficking directs constitutive cargo secretion at distinct sites of the plasma membrane, cilia, and junctional structures. Central to this machinery is the exocyst complex, which tethers cargo vesicles to their destination membrane, alongside other intracellular membrane tethering roles. Precisely how the exocyst spatially integrates membranes and membrane resident binding partners is unclear. Here, we address the structural morphology and formation of the human exocyst complex. Through structural approaches coupled to predictive models, we determined that the exocyst and its subcomplexes have extended ‘arm-like’ structures that help maximize its reach. Moreover, we demonstrate minimal intersubunit interaction, in contrast to prior models. Nucleation of the holocomplex occurs through a single site, explaining its spatial extension. Our results provide the biochemical basis for exocyst complex assembly, suggesting an ornate extended architecture.

**Significance:** Cargo transport to the eukaryotic cell plasma membrane predominantly relies on the exocyst as a central, signal-integrating polarized trafficking complex. How this single cargo vesicle tethering complex can target diverse vesicles to destination membrane is poorly understood, partly due to inconsistent structural models. Here, we show an extended architecture of the human exocyst complex, revealing a minimal nucleation interface between its two subcomplexes. The distinct morphology of human exocyst, compared to yeast models, suggests a novel mechanism that supports its versatile role in membrane trafficking.

## Introduction

Metazoa have vastly expanded trafficking pathways. This increase in complexity compared to other eukaryotes is coordinated by the versatile cellular machinery that controls these pathways. One such machinery delivers cargo to the plasma membrane and specialized structures of the cell such as cilia^1–3^. In this polarized trafficking pathway, the heterooctameric exocyst complex takes a central role^2,3^. Cargo include those constitutively secreted, but increasing evidence implicates the exocyst complex in exosome^4^, GLUT4^5,6^, and lysosome secretion^7^, cilia recycling^8^, and autophagy^9^. Thus, flexible adaptability of the exocyst is suggested to enable function amongst biophysically distinct cargo vesicles^10^. How this versatility could be achieved is not well understood.

The complexity of exocyst-mediated cargo capture is apparent in the large number of putative regulatory partners, including actin remodelers^11^, coiled-coil tethers^12^, SNARE proteins^13^, a number of small GTPases^3^, motor proteins such as myosin 5^14^, and phosphoinositide signaling lipids^15,16^. In contrast, coiled-coil tethers typically have few binding partners^17^. Despite a structural model of the yeast exocyst, the mechanistic integration of exocyst binding partners both structurally and functionally is poorly understood^18^. Yet, such a functional model is required for targeted modulation of the exocyst across its diverse roles.

The octameric exocyst is a member of the Complexes Associated with Tethering Containing Helical Rods (CATCHR) family, which all contain homologous subunits evolved from a common, monomeric ancestor^17,19–21^. Tethering complexes specifically initiate cargo delivery through direct capture of a cargo vesicle at its destination membrane^21^. This capture requires a recognition of membrane identity, ultimately through binding to small GTPases and phosphoinositides^22–25^. Owing in part to biochemical intractability, we lack mechanisms and structural models for how the exocyst, or indeed other CATCHR family members, capture cargo.

Interestingly, structural models of the yeast exocyst reveal a largely constrained and ellipsoidal complex^26–28^, an exception to other CATCHR family architectures^29,30^. These models and biochemical approaches reveal structure revealed two key multiprotein assemblies. Both Subcomplex-1 (mammalian Exoc1:Exoc2:Exoc3:Exoc4; yeast Sec3:Sec5:Sec6:Sec8, respectively) and Subcomplex-2 (mammalian Exoc5:Exoc6:Exoc7:Exoc8; yeast Sec10:Sec15:Exo70:Exo84, respectively) assemble in an ordered and anti-parallel manner via an N-terminal coiled-coil contributed by each subunits^27,31^. Following these coiled-coils is an approximately 15 nanometer C-terminal α-helical rod, generally extending away from the core coiled-coil of each subcomplex. In previous yeast structural models, the two subcomplexes clasp together forming a constrained architecture. The C-terminal α-helical domains are generally bundled together and pointing in the same direction. Thus, presumably extensive intersubunit interfaces are buried, as observed in the CATCHR family member Dsl1^26,28,32^. Besides the well-defined coiled-coils, the remaining exocyst interfaces, and particularly those responsible for holocomplex formation, are not defined.

In contrast to electron microscopy approaches, combinatorial fluorescent-tagging of the yeast exocyst followed by *in vivo* microscopy revealed an overall open-handed architecture^33^. Moreover, studies of functional cargo delivery suggest both disassociation of exocyst subcomplexes and even individual subunits^34–36^. The reconciliation of a restricted, constrained complex with both disassembly and structural extension previously observed in plants and human exocyst is challenging in the absence of more structural information^37,38^.

Binding partner recruitment may result in increased conformational extension of the exocyst. Allosteric regulation of the exocyst by the Rho/Cdc42 small GTPases results in increased binding to the SNARE membrane fusion machinery^39^. Simultaneously, activation by the beta-propeller Sro7 appears to unleash a distinct conformational change, resulting in increased Sec4 (mammalian Rab11) binding and tethering^40^. In contrast to these examples of increased exocyst extension, the CATCHR family member Dsl1 adopts a more constrained architecture upon SNARE binding^32^. Despite this mechanistic structure-function progress, models for mammalian exocyst structure and function are elusive yet required to understand its variably distinct roles in polarized trafficking.

Here, we have reconstituted human exocyst complex formation and characterized the subcomplex and holocomplex structural architecture. We combine predictive structural models, based upon biochemical assembly, and determine an overall structural morphology for the human exocyst. Predictive structural ensembles enabled us to identify a key nucleating feature for holocomplex formation from subcomplexes. These data reveal the extended architectural framework of the human exocyst, explaining its versatile mechanism across diverse cargo and functional roles.

## Results

### Human exocyst subcomplexes have minimal intersubunit interactions and are stable

The exocyst complex is formed from two subcomplexes consisting of subunits Exoc1-4 (Subcomplex-1) and Exoc5-8 (Subcomplex-2) which assemble through N-terminal coiled-coils. Moreover, a key linkage is formed between Exoc4 and Exoc5^25^. However, the precise intersubunit interfaces are unclear^25,27,28,31,33,41^. To obtain a guiding structural model for exocyst subcomplexes, we assembled a structural model derived from human hierarchal assembly experiments (Fig. 1A,B; Fig. S1A-C). Analogous to the yeast complex, the single subunits assemble at their N-terminal coiled-coils, resulting in splayed subcomplex models, reminiscent in form to CATCHR family members. Moreover, direct predictions with AlphaFold Server including all templates (i.e. PDBID: 5YFP^28^) resulted in similar, yet somewhat more constrained structural models (Fig. S1D-G). Importantly, we observe minimal intersubunit interactions in the C-terminal α-helical repeats.

**Figure 1.**
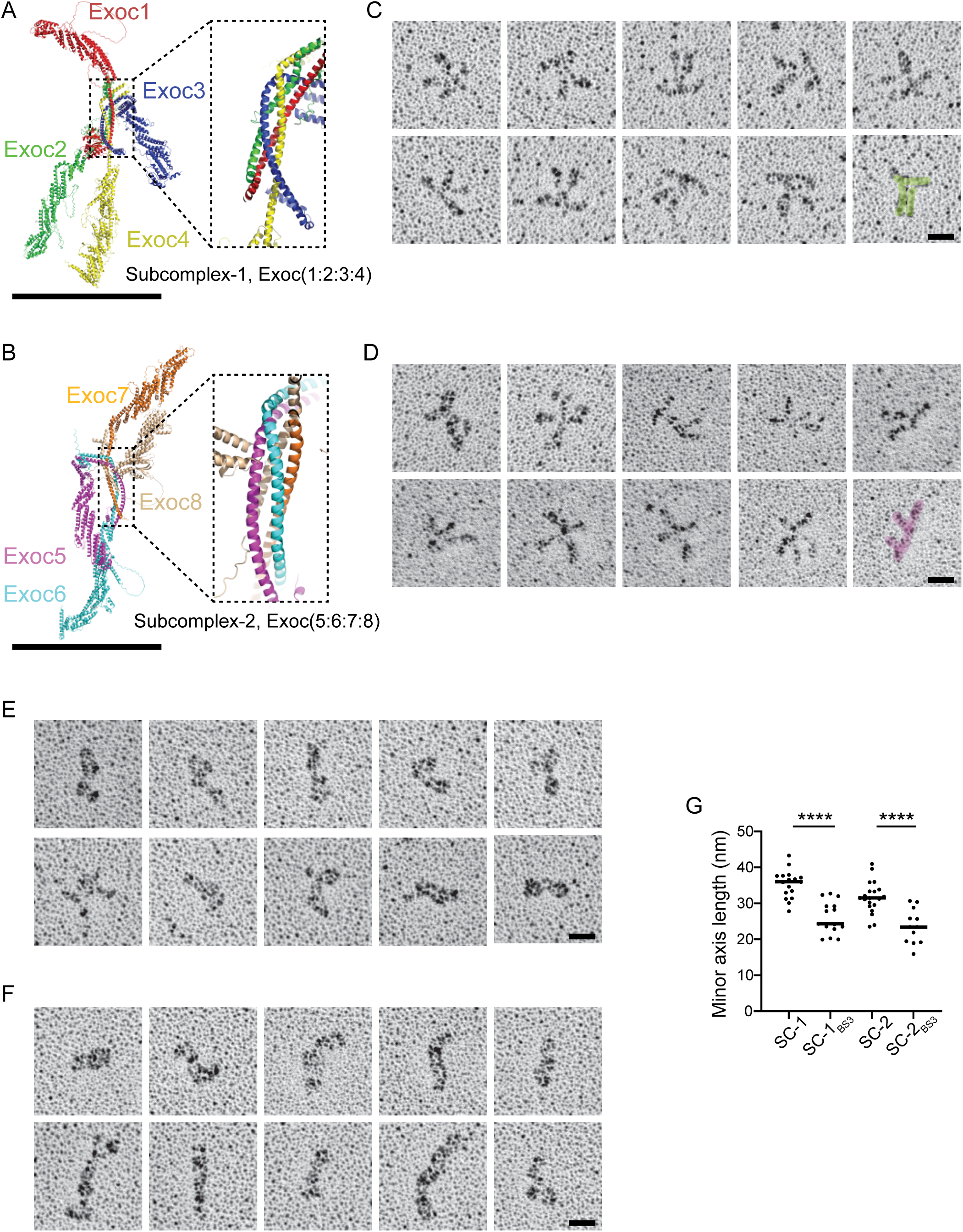
Exocyst subcomplexes are extended but constrained upon crosslinking. **A**, Structural architecture of human exocyst Subcomplex-1 derived from AlphaFold2 modeling. Subunits Exoc1 in red; Exoc2 in green; Exoc3 in blue; Exoc4 in yellow. N-terminal coiled-coil highlighted. Scale bar, 20 nm. **B**, Structural architecture of human exocyst Subcomplex-2 derived from AlphaFold2 modeling. Subunits Exoc5 in magenta; Exoc6 in cyan; Exoc7 in orange; Exoc8 in wheat. N-terminal coiled-coil highlighted. Scale bar, 20 nm. **C**, Representative examples of rotary-shadowing electron microscopy of human exocyst Subcomplex-1, n=18, and **D**, Subcomplex-2, n=20. Bottom right panel shows particle highlighted in green (C) and magenta (D). Scale bar, 20 nm. **E**, Representative examples of rotary-shadowing electron microscopy of BS^3^ crosslinked and repurified Subcomplex-1, n=15, and **F**, Subcomplex-2, n=12. Scale bar, 20 nm. **G**, Measure of minor axis length for native and crosslinked Subcomplex-1 (SC-1) and Subcomplex-2 (SC-2). P<0.0001, ****.

Structural models of yeast exocyst, supported by cross-linking mass spectrometry, suggest intersubunit interactions with no yet defined interfaces^26,28^. In contrast, other membrane tethering complexes largely extend, positioning regulatory interactions to the periphery^29,30,42,43^. To experimentally define structural extension in human exocyst, we purified Subcomplex-1 and Subcomplex-2 by multi-gene baculovirus-mediated insect cell expression (Fig. S2A-D)^44^. Throughout purification, we analyzed small-volume fractions by mass photometry^45^. This approach enabled precise identification of protein fractions meeting theoretical molecular mass criteria. Importantly, we observed no aggregated material or disassembly, providing strong evidence of the biochemical homogeneity and stability of both subcomplexes.

### Exocyst subcomplexes are entirely extended with four clear arms

To experimentally validate the structural predictions demonstrating subunit extension, we examined exocyst subcomplexes by platinum rotary shadowing electron microscopy to define their overall two-dimensional structural architecture^18,46^. This approach applied to Subcomplex-1 and Subcomplex-2 revealed architectures surprisingly consistent with AlphaFold models (Fig. 1C,D). Both subcomplexes adopt an extended cross shape, with occasional views illuminating a potential transition from an open ‘X’-to a more closed ‘T’-shaped conformation. Thus, the arms defined by the C-terminal α-helical repeats of the exocyst subunits, extend outwards with no observable, consistent intersubunit interaction in this region.

Prior structural approaches used crosslinking or fixation both for biochemical purposes and to provide additional structural restraints^26,28^. To ascertain the impact of crosslinking to exocyst structural architecture, we crosslinked and repurified subcomplexes by size-exclusion chromatography (Fig. S2E). Crosslinked subcomplexes remained well-behaved in size-exclusion (Fig. S2E) and in mass photometry analysis (Fig. S2F). Next, we analyzed amide-linked samples by rotary shadowing. Subcomplex particles appeared significantly different from their native state (Fig. 1E, F), appearing constrained, and more consistently so on Subcomplex-2 than Subcomplex-1. Unlike in native conditions (Fig. 1C,D), the C-terminal α-helices of the subunits could not be distinguished. We quantified the spread of the subcomplex by measuring the constraining lengths along the minor particle axis, revealing a significant reduction for crosslinked subcomplex (Fig. 1G), and demonstrating their restricted conformation when crosslinked.

### Exocyst holocomplex adopts an extended conformation

The extension of exocyst subcomplexes suggested that their assembly, dependent on an Exoc4:Exoc5 interface^25,27^, may result in a consolidation of C-terminal α-helical repeats, or potentially an even more extended structure^37^. To experimentally resolve this question, we purified exocyst holocomplex and validated its monodispersity by sizing, mass photometry, and light scattering (Fig. S3A-C). We applied this material to rotary shadowing electron microscopy and observe large, extended complexes (Fig. 2A). These exocyst holocomplex particles reveal a morphology reminiscent of linking subcomplexes (Fig. 1C,D) through each contributing a single subunit. Occasionally, subparticle investigation reveals a view remarkably similar to that observed in the subcomplex analysis (Fig. 1C,D; Fig. 2A, shaded in lower right panels). We observed that the C-terminal α-helices of the subunits are not generally restricted, suggesting an extended human exocyst architecture.

**Figure 2.**
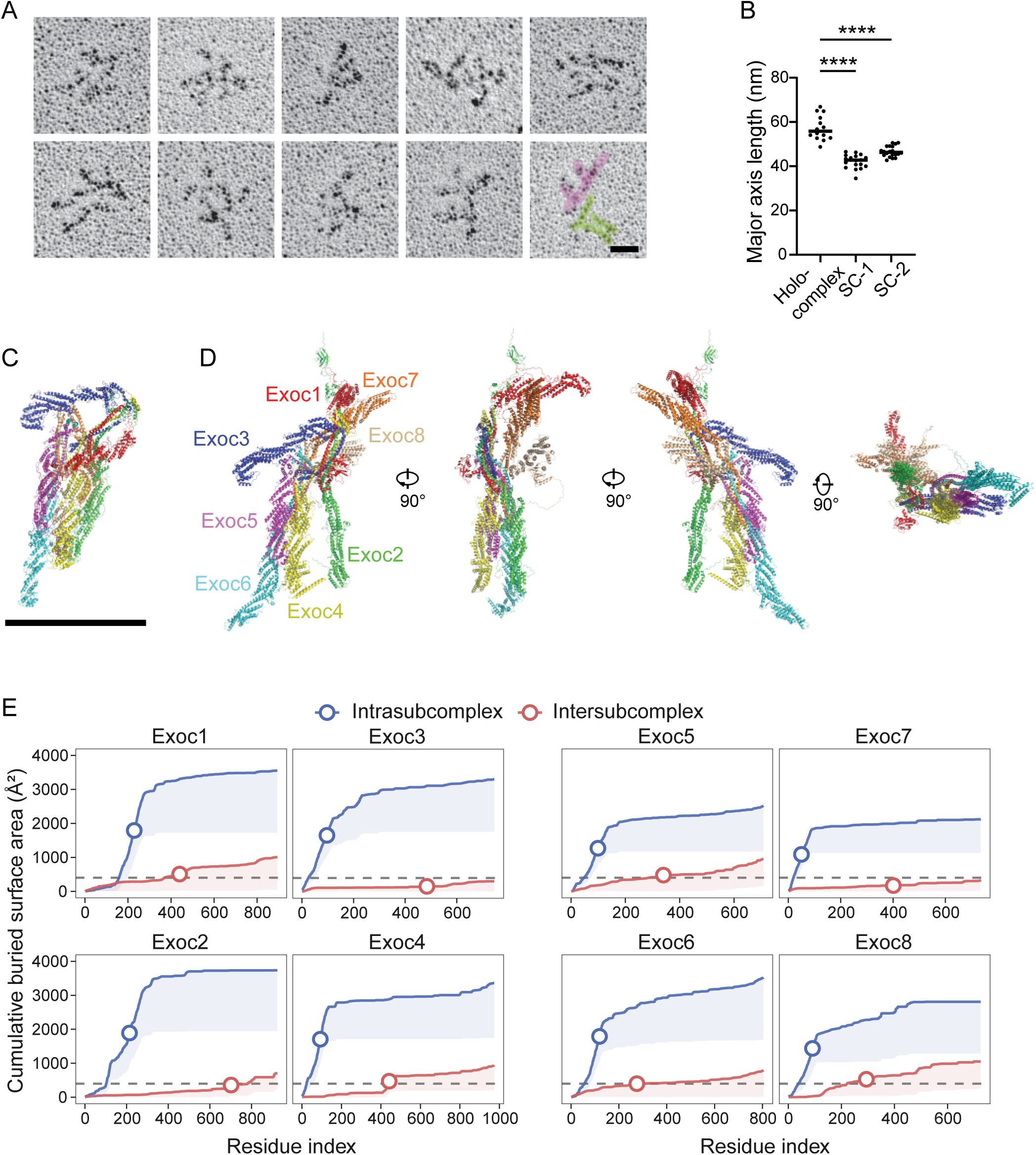
Exocyst holocomplex is extended with minimal intersubcomplex interactions. **A**, Representative examples of rotary-shadowing electron microscopy of human exocyst holocomplex, n=14. Bottom right panel shows particle highlighted in green and pink. Scale bar, 20 nm. **B**, Measure of major axis length for exocyst holocomplex and Subcomplex-1 (SC-1) and Subcomplex-2 (SC-2). P<0.0001, ****. **C**, Representative yeast exocyst structure (cryo-EM model, PDB ID: 5YFP). Scale bar, 20 nm. **D**, Structural architecture of human exocyst holocomplex derived from AlphaFold2 modeling, same scaling as **C**. **E**, Structural ensemble analysis. Plots of subunit amino acid residue index versus cumulative buried surface area for intrasubcomplex (blue) and intersubcomplex (red) interactions. Circles represent midpoint of maximal area. Shaded areas represent the cumulative 95% lower confidence internal of the mean buried surface area. Dash lines are at 400 Å², which is a cutoff for stable interfaces.

Importantly, we observe only minimal indication of intersubunit interactions in the C-terminal regions. To quantify exocyst size, we measured the length along the major axis (Fig. 2B). The measured distances suggest a broad extension upwards of 50 nm and demonstrates the extension of the human exocyst holocomplex. However, this observation is in stark contrast to prior yeast structures. Despite extensive screening we have been unable to resolve high-resolution maps by cryo-electron microscopy. However, we have observed human exocyst by negative-stain electron microscopy, which reveals unaggregated and ellipsoidal particle classes consistent with previous observations by this approach (Fig. S4).

### Exocyst models suggest intrasubcomplex interfaces predominate

With no biochemical evidence of large interfaces between the exocyst subcomplexes or individual subunits^25^, the exocyst assumes a far more extended conformation than previously appreciated. Indeed, early studies on both human and plant exocyst structure suggested extended subunits^37,38^, in sharp contrast to that observed in yeast models (Fig. 2C). To structurally explain the extended architectures we observed, we generated an exocyst holocomplex model based on AlphaFold prediction and subsequent refinement. In this model, the C-termini were similarly extended, with minimal intersubunit interactions (Fig. 2D).

To quantify the predicted extension, we modelled in AlphaFold2-multimer (AF2-M) the human exocyst complex excluding the yeast structural templates (Fig. S5A). We repeated this prediction numerous cycles with unique seeds to create a structural ensemble. Indeed, in all predictions the coiled-coil regions consistently assemble, confirming their anti-parallel orientation and the overall topology of the subcomplexes^28^. Strikingly, the models are greatly variable in their spatial orientations, and no specific interfaces are modelled between the alpha-helical repeats except for the consistent Exoc4:Exoc5 link, validating prior data suggesting this interface^25^.

Because of morphological variability in the models, we quantified the overall difference in intra- and inter-subcomplex interaction through measuring the buried surface area for each exocyst subunit. This structural interface analysis reveals that the inter-subcomplex interaction surfaces of individual subunits are relatively small compared to intra-subcomplex interactions (Fig. 2E). Moreover, in all intrasubcomplex interactions, we observe a large increase in the buried surface area of the N-terminal coiled-coil bundles, likely forming during cotranslation^47^.

Moreover, an Exoc4:Exoc5 interface is apparent, demonstrated by the burst of buried sequence after Exoc4 residue 400 (interface midpoint at residue 444; Fig. 2E). Other than this subunit pair, inter-subcomplex interactions are minimal. In these human exocyst predictions, the interface sizes of intra- and inter-subcomplex differ significantly, observed by calculating the average interface size across all subunits and predictions (Fig. S5B). Interestingly, AF2-M models of yeast exocyst shared similar characteristics to human (Fig. S5C). Thus, a predictive approach evidences that the intrasubcomplex interfaces, defined by the N-terminal coiled-coil regions, largely predominate over intersubcomplex interfaces.

### Spontaneous assembly of exocyst holocomplex

Exocyst assembly requires the coalescence of the two subcomplexes (Fig. 3A) and may be regulated^48^. To test this assembly *in vitro*, we combined purified subcomplexes and assessed their binding. An estimated K_D_ of 78 nM (Fig. S5D) enabled us to harness mass photometry to evaluate complex formation^49^. We preincubated independently purified Subcomplex-1 (353.4 kDa; theoretical 404.0 kDa) and Subcomplex-2 (313.3 kDa; theoretical 337.7 kDa) at equal concentration, and measured the subsequent mass of the potential complex. Near complete (74%) assembly of the exocyst holocomplex (695.7 kDa; theoretical 741.7 kDa) was observed (Fig. 3B). Thus, exocyst subcomplex assembly occurs in the absence of binding partners to form the stable holocomplex.

**Figure 3.**
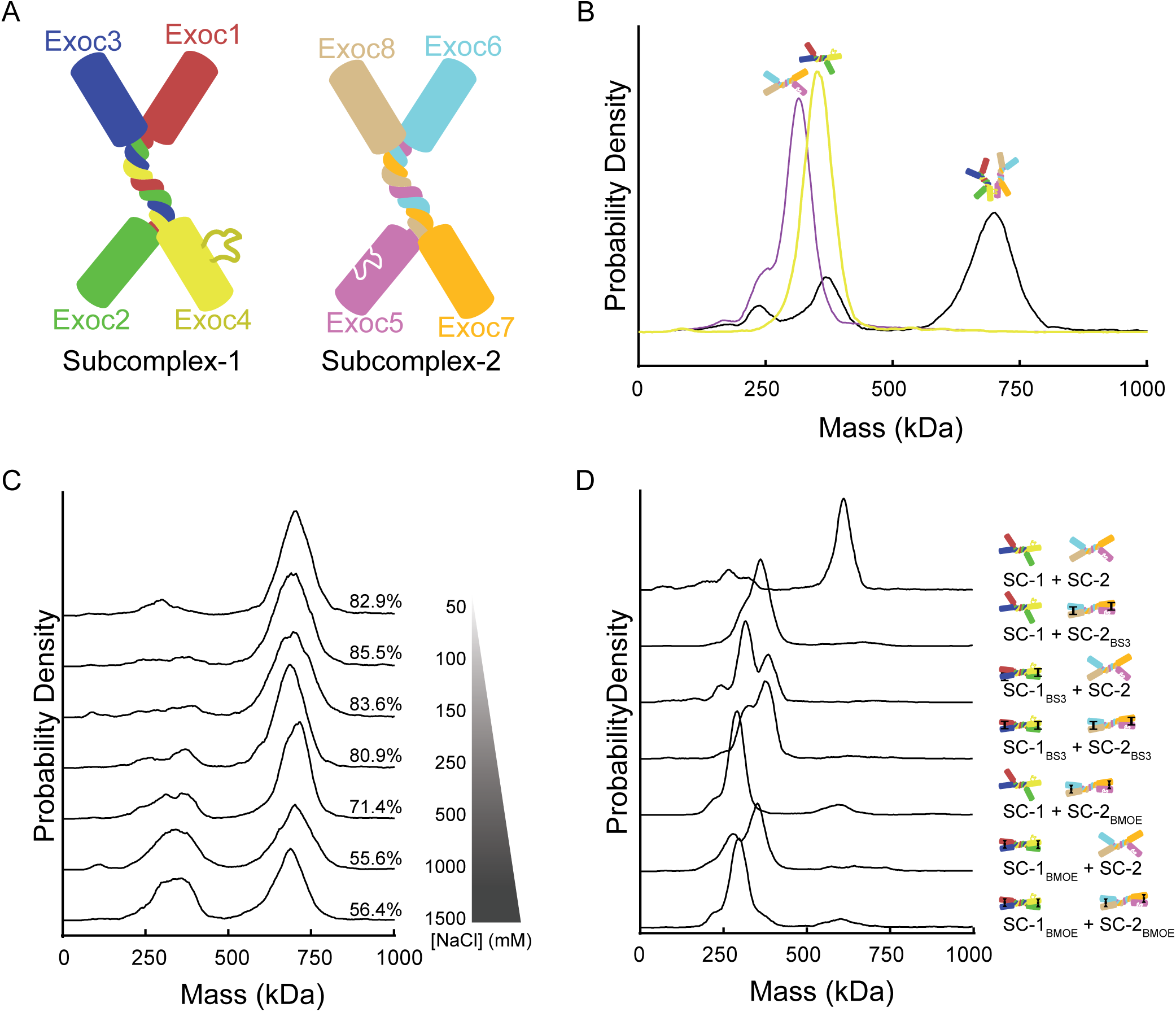
Subcomplex coalescence to holocomplex, and its abrogation by restriction. **A**, Schematic model of exocyst subcomplexes, colored as in all predictions. Exoc4:Exoc5 connection hypothesized as loop-pocket on subunits. **B**, Mass photometry plot for individual subcomplexes and their combination. Representative experiment, n=6. **C**, Mass photometry plots for holocomplex coalescence in varying salt conditions. Representative experiment, n=3. **D**, Mass photometry plots for subcomplex coalescence under BS^3^ and BMOE crosslinking conditions. Note absence of higher-order aggregates. Schematics describe experimental condition. Representative experiment, n=3.

The formation of large protein complexes most typically occurs cotranslationally^50^, with assembly burying large hydrophobic surfaces. Because the subcomplexes were well behaved but could spontaneously coalesce to form holocomplex, we evaluated the role of electrostatics in assembly by varying salt conditions^51^. Holocomplex formation was decreased proportionally with increased salt, reaching a minimum at the theoretical Debye limit (Fig. 3C)^52^. Thus, electrostatics have a significant role in exocyst holocomplex coalescence, which is remarkably stable *in vitro* in the absence of further regulation.

Structural studies of yeast exocyst strongly suggest large interfaces between exocyst subcomplexes^26,28^. Thus, we reasoned that restriction of conformation for an individual subcomplex would prevent its incorporation into the holocomplex. To test this hypothesis, we cross-linked and repurified Subcomplex-1 and Subcomplex-2 with both BS^3^ amide- and BMOE cysteine-targeting reagents (Fig. S2E). We qualified their monodispersity by mass photometry (Fig. S2F) and observed no significant difference from native protein in this approach. Indeed, exocyst holocomplex assembly failed when either subcomplex was restricted with BS^3^ or BMOE (Fig. 3D). Thus, the exocyst complex coalescence is abrogated in crosslinking conditions.

### Identification of a conserved interaction site between exocyst Subcomplex-1 and -2

The structural details of exocyst inter-subcomplex interaction are poorly understood, yet essential to understand their function in tethering and regulation by binding partners^39,40^. To model the interactions between Subcomplex-1 and Subcomplex-2, we assembled structure predictions of Exoc4 with Exoc5 (Fig. 4A), focusing only on the C-terminal α-helical repeats. Indeed, these α-helical repeats were highly modellable (Fig. S6A-C).

**Figure 4.**
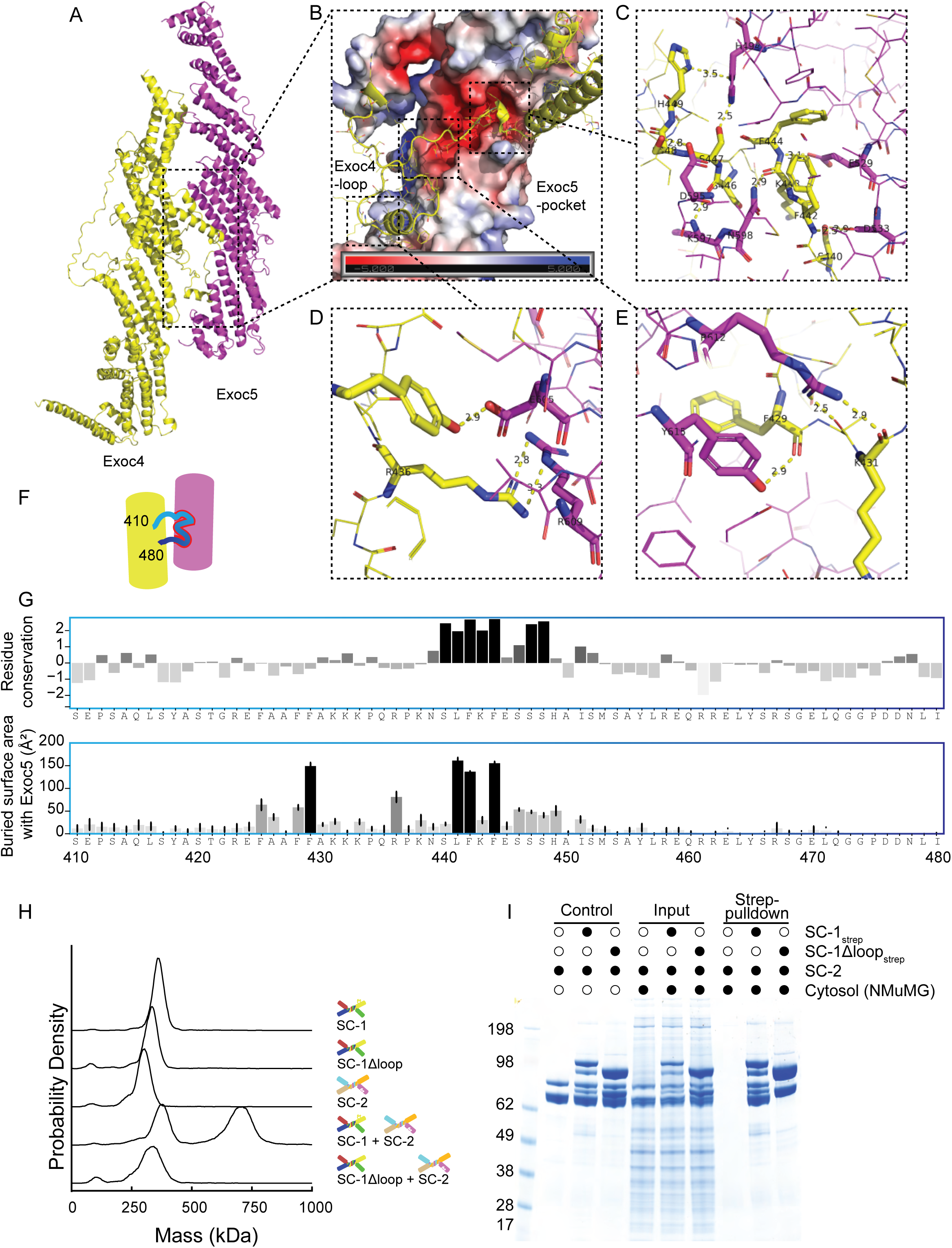
A conserved loop-pocket interaction drives exocyst holocomplex formation. **A**, Interaction prediction of C-terminal domains of Exoc4 and Exoc5 by AlphaFold2. C-terminal Exoc4 is colored yellow, C-terminal Exoc5 colored magenta. Extended loop highlighted. **B**, Zoom of the Exoc4:Exoc5 connection. The loop domain of Exoc4 settles in the negatively charged pocket of Exoc5. **C-E**, Electrostatic and hydrogen bonds are shown in yellow dash lines. The relevant amino acids are shown in bold sticks. **F**, Schematic of Exoc4:Exoc5 loop interaction with loop colored as in panel G. **G**, Residue conservation (top) and the average residue-level buried surface area of the Exoc4 loop formed with Exoc5, across the structural ensembles (bottom). **H**, Mass photometry plots for holocomplex coalescence from subcomplexes, comparing SC-1 and Subcomplex-1Δloop (SC-1Δloop). Representative experiment, n=3. **I**, Subcomplex pulldowns analyzed by SDS-PAGE. Pulldown of SC-2 by SC-1 variants to measure holocomplex coalescence from subcomplexes. Comparing SC-1 and SC-1Δloop. Representative experiment, n=2.

Intriguingly, in Exoc4:Exoc5 models, we identified a single, electrostatically positive loop (T410-V492) reaching outward from the alpha-helical repeat region of Exoc4 (Fig. 4A). In the prediction, this loop interacts with a pocket lined with complementary negative charges in Exoc5 (Fig. 4B). The predicted interaction region contained 15 hydrogen bonds between the Exoc4 loop and Exoc5 pocket (Fig. 4C-E), centered around a highly conserved phenylalanine-442. Indeed, a significant portion of this loop appears constrained in healthy populations, ranking within the 25th percentile of the most missense-intolerant sites in Exoc4^53^. Moreover, across the ensemble AF2-M predictions we assembled (Fig. S5A), we noted a high degree of amino acid conservation (particularly surrounding the central phenylalanine), and significant buried surface area in the loop region (Fig. 4F,G). Strikingly, this predicted Exoc4:Exoc5 loop-pocket interaction suggests a single, predominant interaction nucleating holocomplex coalescence across multiple species (Fig. S6D,E).

### Breaking the exocyst through Exoc4 loop deletion

To test the requirement for Exoc4 T410-V492 in human exocyst complex formation, we expressed and purified a Subcomplex-1ΔLoop complex variant (Fig. S7A). We evaluated holocomplex assembly by mass photometry in the presence of excess Subcomplex-1 and Subcomplex-1ΔLoop. Indeed, the simple deletion of Exoc4 T410-V492 resulted in abrogated exocyst assembly (Fig. 4H). Moreover, we coexpressed GST-fusions of Exoc4 and Exoc4ΔLoop with each individual exocyst subunit (Exoc1-8), and confirmed that the deletion mutant maintained the Exoc1 and Exoc3 binding previously observed (Fig. S7B,C)^25^. Thus, human exocyst holocomplex formation specifically requires an interaction between Exoc4 and Exoc5, nucleated by a single loop.

Exocyst assembly in cells may utilize other factors^48,54^. Therefore, we reasoned that cellular factors may overcome an assembly requirement for the Exoc4 loop in forming holocomplex. To test for factors enabling the interaction between the two subcomplexes, we incubated cytosolic extracts with purified subcomplexes followed by a pulldown assay (Fig. 4I). Subcomplex-2 was captured in the presence of Subcomplex-1, but not in the presence of Subcomplex-1ΔLoop. Hence, exocyst holocomplex formation is broken in the context of a single loop deletion in Exoc4, irrecoverable by inclusion of cell factors.

## Discussion

Structural models of the human exocyst have proven elusive. Here, we aimed to determine a structural model that explains the varied and diverse functions of this complex in metazoan cell biology. Structural prediction coupled to electron microscopy and state-of-the-art biochemistry approaches enabled us to define an extended exocyst morphology. The outward reaching arms of the exocyst are enabled by coalescence of its two constitutive subcomplexes at a single site. These results form a foundation for cell biological exploration of exocyst structure-function.

Generally, CATCHR complex subunits form tight N-terminal coiled-coil bundles, explicitly shown for exocyst, COG, and Dsl1^28,55,56^. This coiled-coil is followed by extended helical rods, which form the bulk of known interaction sites. Both the GARP and COG complex share a similar hub-and-spoke assembly, with the coiled-coil bundle forming the core. Thus, this subunit configuration of CATCHRs may be stabilized or take on restricted orientations through binding partners, as in the Dsl1 complex^32^. Here, we have demonstrated that the human exocyst also takes on an extended hub-and-spoke assembly, spanning up to 60 nm in space along its longest axis (Fig. 5A). This breadth is enabled by holocomplex formation derived from just a pair of subunits connected by a tiny proportion of the total amino acids. Such a small connection may also support a relatively large twist of Exoc4 relative to Exoc5 (Fig. 5B). Indeed, heterogeneity in this interaction was observed in the AlphaFold3 structural ensembles. While the experimental limitations of the rotary shadowing approach limit us only speculate on the three-dimensional molecular-scale structure of the complex, predictive models provide a further conceptual means to contemplate mechanistic function.

**Figure 5.**
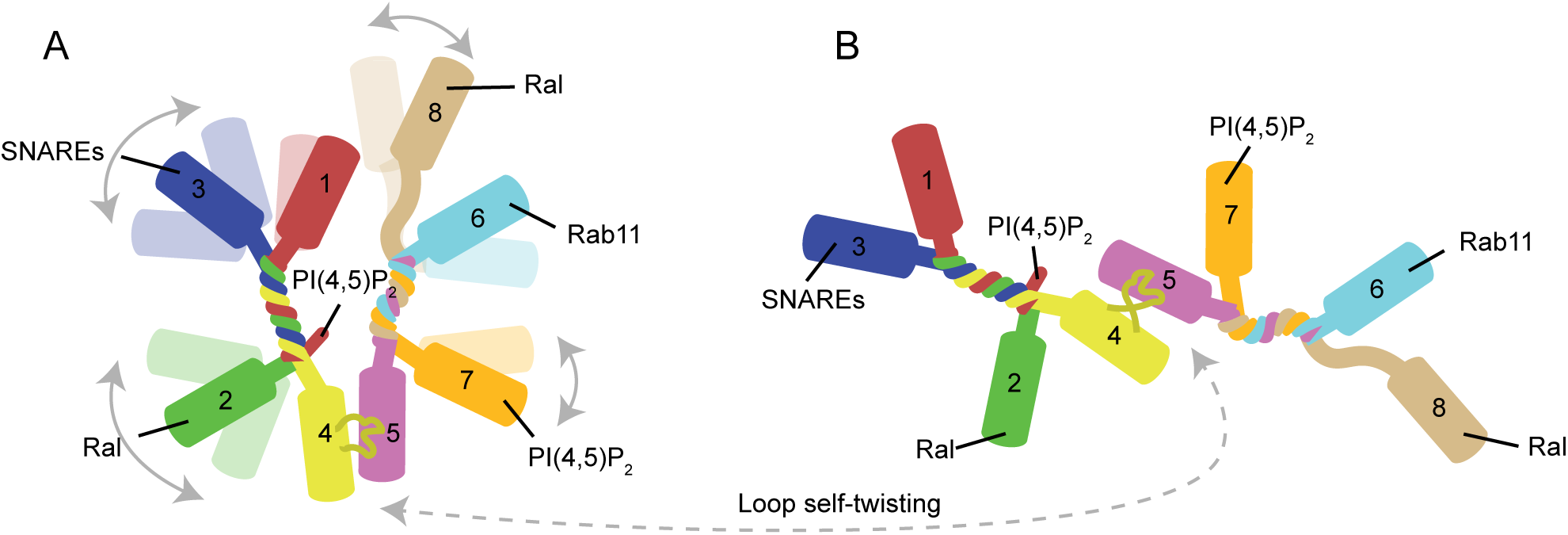
Schematic concept of exocyst structure. **A,** Schematic model of exocyst holocomplex, colored as in all predictions. The extended alpha-helical rods of each subunit extend outwards from the inner core formed from the coiled-coil regions for each subcomplex, and the Exoc4:Exoc5 connection linking the two subcomplexes. The extended rods each harbor distinct binding partners previously biochemically verified, with an obvious potential for flexibility in such an architecture. **B**, The extension of some holocomplexes, and the length of the Exoc4 loop may enable a twist of the subcomplexes with respect to one another, further enabling a flexible adaptation for distinct cargo vesicle recognition.

The three-dimensional extension observed for the C-terminal α-helical rods of individual subunits places the interactions for most known binding partners well into the periphery of the structural architecture. Amongst structurally-interrogated binding partners, that of Rab11 on Exoc6^57^, the signaling lipid phosphatidylinositol (4,5)-bisphosphate (PIP_2_) on Exoc7^15^, and the Ral binding sites on Exoc2 and Exoc8^58,59^ each occur at the C-terminal α-helical rods. Thus, by positioning to the extremity, steric occlusion is entirely unlikely. Indeed, exocyst avidity in cargo vesicle recognition may have a role in programming versatility.

In contrast to most binding partner interaction sites, the PIP_2_ binding PH domain at the N-terminus of Exoc1 (prior to the coiled-coil) has effectively no linker. This situates it directly in the vicinity of the Subcomplex-1 core coiled-coil bundle. In contrast, the yeast homologue Sec3 has an extended linker that enables transient conformations of this lipid binding domain, with occasional forays into the region of Sec6 (Exoc3) in a manner positioning it coincident with the other PIP_2_ binding site on Exo70 (Exoc7)^26^. Thus, fixed PH domain positioning at the core of Subcomplex-1 may enable human exocyst the capacity for *trans*-PIP_2_ binding to drive a mode of tethering, as we previously observed^25^. Higher-resolution models are required to determine these aspects with certainty.

A dilemma for structural-functional study of large and extended protein complexes like exocyst is the reduction of flexibility. The human exocyst holocomplex is more heterogeneous in appearance than previously observed for the yeast exocyst^28^, even in the presence of activating partners^39^. Fixation reagents or crosslinkers generally restrict specimen homogeneity, eliminating potential conformations. Of note, all structural holocomplex studies to date have been performed either in the presence of crosslinkers including uranyl acetate/formate or on substrates^27,28^. Recent work on Dsl1, perhaps the simplest of the CATCHR family members, begins to broaden these approaches by harnessing binding partners to restrict degrees of freedom^32^. Indeed, this concept may pave the way for high-resolution human exocyst models.

Moreover, we speculate that flexibility of the α-helical rods, demonstrated to behave as mechanical springs^60^, enables the versatility in cargo capture. An extended *and* flexible structure will explore more conformation space, has increased likely of ligand/binding partner capture, and generally better explains the cell biological role. However, this hypothesis requires further experiments and remains speculative.

Although the structural-functional relationship of exocyst remains controversial, a more dynamic and flexible framework is favored by the versatility of its function. Indeed, limitations of rotary shadowing and other surface-bound electron microscopy approaches more likely results in an abrogation of conformational heterogeneity. Hence, solution structure approaches will prove critically important to resolving the molecular-scale architecture of this vast complex, defining the molecular basis for its conserved and essential roles in Metazoa.

## Acknowledgements

We thank Nicholus Mukhwana for cytosol extracts and the Murray lab for experimental support. We thank Yogesh Kulathu and Kirby Swatek for careful reading. We personally acknowledge the technical expertise of Marlene Brandstetter and all at the Electron Microscopy Facility at Vienna BioCenter Core Facilities (VBCF), member of the Vienna BioCenter (VBC), Austria. This work has made use of the resources provided by the Edinburgh Compute and Data Facility (ECDF) (http://www.ecdf.ed.ac.uk/).

## Methods

### Plasmid construction

Exocyst subunit plasmids were a kind gift from Channing Der (Addgene #53755-53762). Each was subcloned into FlexiBAC SF9 expression plasmids^61^. Subsequent modifications were performed by standard molecular biology approaches. All plasmids generated in this study will be made available publicly on Addgene.

### Protein expression

Exocyst subunits, subcomplexes, holocomplex and relevant variants were all expressed in SF9 cells (Oxford Expression Technologies) via FlexiBAC baculovirus expression system^61^. Briefly, to generate a baculovirus, SF9 cells at 5 × 10^5^ cells/mL were co-transfected by the plasmid harboring the gene(s) of interest and the linearized DefBac viral backbone using ESCORT-IV transfection reagent (Sigma) in a 6-well suspension plate (ThermoFisher). After 5 days of incubation, cells were checked for signs of transfection – a successful transfection enlarged the cells from ~Ø12nm to ~Ø19nm. 50-100 μL of the supernatant (P1 virus) was used to transfect 50 mL of SF9 cells at 5 × 10^5^ cells/mL for the P2 virus. After another 5 days of incubation, cells were checked, and the supernatant (P2 virus) was ready for large scale protein expression.

For exocyst subunit and subcomplex expressions where only single specific viruses were required, 5-10 mL of the relevant P2 viruses were used to infect 1L of SF9 cells at 1 × 10^6^ cells/mL, respectively. For exocyst holocomplex expression, 1L of SF9 cells at 1 × 10^6^ cells/mL were co-infected with 4 mL of Twin-Strep-tagged Subcomplex-1 and 6 mL of untagged Subcomplex-2. A relative redundance of untagged Subcomplex-2 during the co-expression assured the proper stoichiometry between the two subcomplexes during further purification steps. After 48-60 hours of incubation, the cells were harvested by centrifuging at 500 × g for 10 minutes. The harvested cell pellet was resuspended with the standard buffer (20 mM HEPES, pH 7.4, 250 mM NaCl, 0.5 mM TCEP, 0.05% CHAPS), flash frozen in liquid nitrogen and subsequently stored at −80 °C.

### Protein purification

Exocyst subcomplexes and holocomplex were purified via a standard purification route including Twin-Strep affinity chromatography, elution/on-bead cleavage, anion exchange chromatography and size exclusion chromatography. Frozen cell resuspension was thawed in water bath at room temperature. Protease inhibitor cocktail (Roche) and Benzonase (Merck) were dissolved in the thawed mixture prior to on-ice homogenization in a 40 mL Dounce homogenizer. After centrifuging at 60,000 × g at 4 °C for 30 minutes, the supernatant harboring the target protein was incubated with pre-equilibrated Strep-Tactin XT 4Flow resin (IBA) at 4 °C for 30 minutes. The resin harboring the target protein was washed with 20 CV of standard buffer prior to either elution using 50 mM of biotin or on-bead cleavage using recombinant HRV-3C protease for at least 3 hours at 4 °C according to experimental purposes. The elution/cleaved sample was diluted to less than 100 mM of NaCl and applied to a Mono Q 5/50 GL anion exchange column. The target protein was eluted at around 19-21 mS/cm during a 100 CV of gradient elution from 100mM to 1M of NaCl. The peak fractions were qualified by Mass Photometry and the best monodispersed fraction(s) was further polished through a Superose 6 Increase 10/300 GL size exclusion chromatography column. Finally, the peak fractions were checked via SDS-PAGE and Mass Photometry, aliquoted, flash frozen in liquid nitrogen and stored at −80 °C.

The peak fractions of anion exchange chromatography during exocyst subcomplex purifications were partially reserved for crosslinking, respectively. For BS^3^-crosslinking, the concentration of BS^3^ was 0.2 mM, 50 to 60-fold of the amount of protein. Mixtures were incubated at room temperature for 30 minutes. For BMOE-crosslinking, the protein samples were buffer-exchanged to PBS buffer, pH 7.4, using Zeba Spin desalting column (Thermo) to prevent reoxidation of disulfides. The concentration of BMOE was 50 μM, 10 to 15-fold of the amount of protein. Mixtures were incubated at room temperature for 1 hour. Finally, the incubations were sized and qualified in the same manner.

### Mass Photometry

Analysis by mass photometry analysis was conducted on a One^MP^ photometer (Refeyn) at room temperature as previously described^62^. AcquireMP software (Refeyn) was used for data acquisition. Briefly, in a typical experiment, 9 μL of buffer was pipetted into the gasket wall without touching the cover glass and subsequently focused in a 5 μm × 10 μm field to a stable sharpness value (around 5.5). 1 μL of diluted protein sample (~0.1 μM) was gently pipetted into the focused drop. The counting events were recorded for 60s at a 1kHz frame rate and processed via DiscoverMP software (Refeyn).

### Exocyst subcomplex incubations

For *in vitro* exocyst assembly experiment under gradient salt concentrations, the purified subcomplexes were buffer exchanged to different salt concentrations using Zeba Spin desalting column (Thermo). Subsequently, the subcomplexes were incubated at final concentration of 0.3 μM at room temperature for 30 minutes and further analyzed by mass photometry. For assembly trials of crosslinked samples, the subcomplexes were incubated at final concentration of 0.3 μM, respectively, at room temperature for 30 minutes and further analyzed by mass photometry. For Subcomplex-1 and -1Δloop incubation with Subcomplex-2, excess 0.6 μM of Subcomplex-1 variants were incubated with 0.3 μM of Subcomplex-2, respectively, at room temperature for 30 minutes and further analyzed by mass photometry.

### Subcomplex pulldown

Pulldown assays were conducted using Eppendorf DNA LoBind tubes. The purified Twin-Strep-tagged exocyst Subcomplex-1 and -1Δloop were incubated with 15μL of pre-equilibrated Strap-Tactin XT 4Flow resin in final 100μL, 0.2 μM, respectively, on a rotary shaker (Biopremier) at room temperature for 30 minutes. Each group of resin was washed 3 times with 1 mL of standard buffer and subsequently applied with a final 0.5 μM of purified Subcomplex-2 to 100 μL. 0.5 mL of extracted cytosol matrix from NMuMG cells was added, respectively, prior to another 30 minutes of incubation. Finally, the resin was washed, eluted with SDS-loading buffer and analyzed by SDS-PAGE.

### Cytosol extraction

NMuMG cells were seeded in a 150 mm dish at 33,000 cells/cm^2^ and grown for 72 hours to high confluence. Cells were then harvested on ice by scrapping in 2 ml of cold DPBS. Cells were pelleted at 1,000 x g for two minutes at 4 °C. The supernatant was discarded, and cells were washed once with 1 ml cold KPBS (containing 25 Mm KCL, 100 mM Potassium phosphate, pH 7.2) and pelleted at 1,000 x g for 2 minutes at 4 °C. The pellet was resuspended in 1 ml cold KBPS and lysed with 30 strokes in Dounce homogenizer. The lysed cells were centrifuged at 18,000 x g for 5 minutes at 4 °C. The post-nuclear supernatant was carefully transferred to a new Eppendorf tube on ice, ready to use. The cytosol extract’s protein concentration (~5 mg/ml) was estimated by NanoDrop.

### SPR

SPR analysis was conducted on a Biacore 8K+ (Cytiva) at room temperature with HBS-P+ buffer (containing 10 mM HEPES, pH 7.4, 150 mM NaCl, 0.05% v/v Surfactant P20, Cytiva). Strap-Tactin XT protein was immobilized to a Series S sensor chip CM5 (Cytiva) following the Twin-Strep-tag capture kit manual. 5 μg/mL (12.3 nM) of Twin-Strep-tagged exocyst Subcomplex-1 was then applied to the chip. To set up the single cycle kinetics, exocyst Subcomplex-2 was diluted to 0.74, 2.22, 6.67, 20 and 60 nM in running buffer along with a parallel blank control for background subtraction. Contact and dissociation time were set for 300 seconds and 600 seconds, respectively, under the flow rate of 10 μL/min. Finally, the chip was regenerated 3M MgCl_2_ under default flow setting. The binding affinity *K_D_* was derived by fitting the association *k_a_*and dissociation *k_d_* with 1:1 binding model via Biacore Insight Evaluation software (version 5.0.18.22102, Cytiva).

### Exocyst subcomplex and full complex prediction via AlphaFold2

N-terminal coiled-coil sequences of Exoc1 (186-285), Exoc2 (172-292), Exoc3 (1-268), Exoc6 (1-139), Exoc7 (1-96), and Exoc8 (1-126) and full-length sequences of Exoc4 and Exoc5 were used to generate a skeletal exocyst backbone model via ColabFold version 1.5.2. The predicted full-length subunits were subsequently aligned with their N-terminal coiled-coils in the exocyst backbone model via PyMol (Version 2.5.7, Schrodinger).

### Exoc4:Exoc5 interaction predictions via AlphaFold Server

Human Exoc4(121-974) and Exoc5(127-708) sequences were submitted in pair to AlphaFold Server. Full-length Exoc4 and Exoc5 sequences from non-human species were submitted in pairs, respectively, to AlphaFold Server. C-terminal interactions and APBS electrostatics of C-terminal Exoc5 were visualized via PyMol (Version 2.5.7, Schrodinger). The sequences of Exoc4 loop domains from different species were extracted for multiple sequences alignment (MSA) using T-coffee with default settings^63^. The MSA result was plotted using Jalview (Version 2.11.3.3)^64^.

### AlphaFold2-Multimer on the human exocyst complex for quantitative analysis

Canonical UniProt sequences of EXOC1-8 (Q9NV70, Q96KP1, O60645, Q96A65, O00471, Q8TAG9, Q9UPT5, Q8IYI6) were used for AlphaFold-Multimer modelling^65^. Predictions were performed with LocalColabFold, running ColabFold version 1.5.5^66^, on an NVIDIA A100 GPU with 500GM of RAM. Template use was explicitly disabled, and the -msa-mode flag was set to “mmseqs2_uniref_env” to increase alignment depth. Eighteen models were generated by varying the random seeds and enabling dropout to sample model uncertainty. All other parameters were set to their default values.

### Structural analysis

Solvent accessible surface area was calculated at residue level using FreeSASA 2.1.2^67^. Relative interface location and Arel(bound) were calculated as previously described^68,69^. The proportion of unstructured residues was determined by assigning secondary structure to the models with DSSP 2.2.1^70^ and calculating the fraction of unassigned secondary structure elements for each chain across the models.

### Rotary shadowing EM and data analysis

Rotary shadowing EM was performed as previously described^46^. Briefly, 0.1 mg/mL of samples were 1:1 diluted in Mabuchi spraying buffer, consisting of 200mM ammonium acetate and 60% (v/v) glycerol, pH 7.4. Diluted samples were sprayed via a capillary onto freshly cleaved mica chips. These mica chips were mounted in the high vacuum evaporator (MED 020, Baltec) under 2L×L10^−5^ mbar and dried. Specimens were coated with 0.7nm of platinum (BALTIC, Germany) at an angle of 4-5°, followed by 6-8nm of carbon (Balzers, Liechtenstein) at 90°. Following deposition, the replica was floated off, picked up on 400 mesh Cu/Pd grids (Agar Scientific, UK) and examined via Morgagni 268D TEM (FEI now ThermoFisher) at a high tension of 80 kV with MegaView III CCD camera (Olympus-SIS).

For the particle size analysis, the micrographs were imported to FIJI^71^ and the particles were picked manually by elliptical selection tool. Measurements of major and minor distance were plotted and compared via t-test in Prism 9 (GraphPad software).

### Negative-stain EM

4 μL of 175 nM purified exocyst sample was applied to a glow-discharged, carbon-coated copper grid (400 mesh, Agar scientific) and incubated for 30 seconds. The excess liquid was gently blotted with filter paper, and 4 μL of 1% uranyl formate was immediately applied as a negative stain. After a 30-second incubation with the stain, the grid was blotted again to remove excess liquid and allowed to air dry completely at room temperature for 10 minutes. The grids were screened using a JEOL 1400 Flash TEM operating at 120 kV. Manual data acquisition was performed under a Gatan Rio camera at 40,000x magnification with a pixel size of 1.4 Å. The total electron dose was optimized to 50 e/Å^2^.

71 micrographs were accepted for particle picking after Patch CTF using CryoSPARC 4.5.1 (Structura Biotechnology). To generate 2D classes for template picking, a total of 27,470 particles were initially selected via Blob Picker, with an average of 387 particles per micrograph. The elliptical blobs were set with a minimum particle diameter of 150 Å and a maximum diameter of 450 Å. After several rounds of 2D classifications, 9,280 particles from 38 classes were selected as template. According to the maximum size of the selected particles, particle diameter was set at 350 Å for template picking. After several rounds of 2D classification, 28,443 particles in 49 classes were finalized for 3 classes of ab-initio reconstitution. The representative model was refined to a final resolution of 6.79 Å.

## Supplemental Figure Legends

**Figure S1.**

**A-C**, Quality control plots for hierarchal AlphaFold2 model.

**D, E**, AlphaFold3 predicted model and positional error plot for Subcomplex-1, and

**F, G**, Subcomplex-2. Scale bar, 20 nm.

**Figure S2.**

**A**, Plasmid construct design for purification of Subcomplex-1 and

**B**, Subcomplex-2.

**C**, Representative purification for Subcomplex-1. Affinity (not shown) cleavage product is purified by anion exchange incorporating fraction collection at ~250 μL volume, with each fraction analyzed by mass photometry. Best fractions representing theoretical mass were taken, combined, and applied to size-exclusion chromatography. Small volume fractions were again taken and analyzed by mass photometry. Best fractions representing theoretical mass were taken and used for experiments.

**D**, Representative purification for Subcomplex-2, as in **C**.

**E**, Size-exclusion chromatograms for Subcomplex-1 (SC-1) and Subcomplex-2 (SC-2), including repurification of crosslinked variants.

**F**, Mass photometry plots for individual subcomplexes used in crosslinking holocomplex coalescence experiments.

**Figure S3.**

**A**, Representative purification for exocyst holocomplex. Affinity (not shown) cleavage product is purified by anion exchange incorporating fraction collection at ~250 μL volume, with each fraction analyzed by mass photometry. Best fractions representing theoretical mass were taken, combined, and applied to size-exclusion chromatography. Small volume fractions were again taken and analyzed by mass photometry. Best fractions representing theoretical mass were taken and used for experiments.

**B**, Thermal denaturation curve for exocyst holocomplex.

**C**, Light scattering plot for exocyst holocomplex.

**Figure S4.**

**A**, Representative images of raw data. Scale bar 500 Å.

**B**, 2D classification of 28,968 picked particles from dataset represented in **A**. **C**, Representative refined Ab-initio reconstitution model.

**D**, Fourier shell correlation shows a refined model resolution of 6.79 Å in **C**. **E**, Viewing direction distribution.

**Figure S5.**

**A**, Representative examples from AlphaFold2-multimer ensemble of models colored as described in Figure 2.

**B**, Interface size and relative position of interfaces for intrasubcomplex and intersubcomplex interactions of human exocyst and

**C**, Yeast exocyst. Box-whisker plots.

**D**, Surface-plasmon resonance measure of Subcomplex-2 affinity for Subcomplex-1.

**Figure S6.**

**A-C**, Exoc4:Exoc5 AlphaFold model quality control plots.

**D**, Sequence alignment of loop region of Exoc4 for several common lab species.

**E**, Interaction prediction of C-terminal domains of Exoc4 and Exoc5 by AlphaFold2 for several species.

**Figure S7.**

**A**, Representative purification for Subcomplex-1Δloop. Affinity (not shown) cleavage product is purified by anion exchange incorporating fraction collection at ~250 μL volume, with each fraction analyzed by mass photometry. Best fractions representing theoretical mass were taken, combined, and applied to size-exclusion chromatography. Small volume fractions were again taken and analyzed by mass photometry. Best fractions representing theoretical mass were taken and used for experiments.

**B,C**, Exocyst subunit assembly experiment analyzed by SDS-PAGE. Pulldown approach for Exoc4 (B) and Exoc4Δloop (C) versus each of eight exocyst subunits. Exoc1, Exoc3, and Exoc5 interactions labeled directly with asterisk.

