## Supplementary figures and images for "Structural extension of the human exocyst is enabled by a minimal interface"

### Supplemental Figures

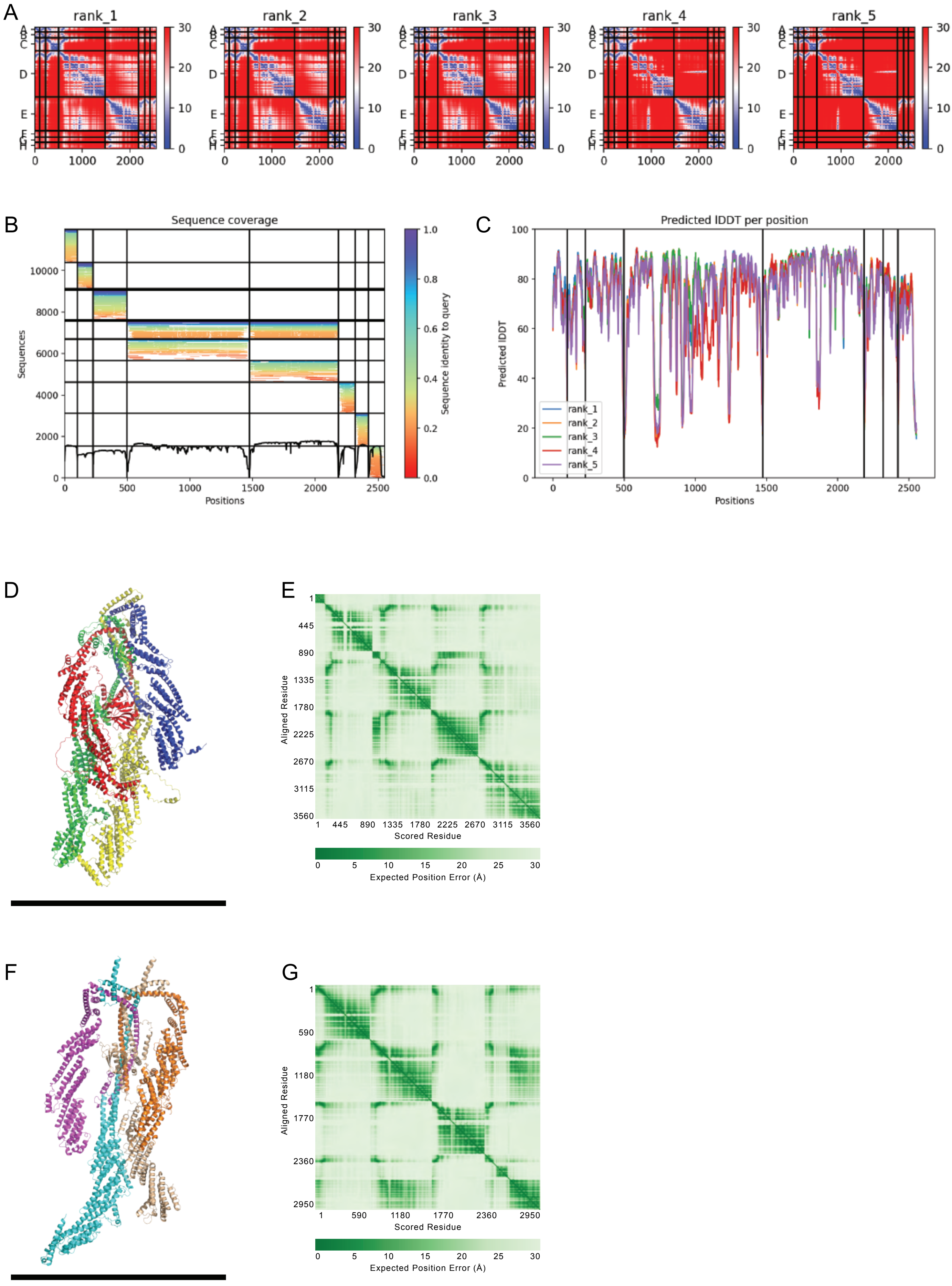

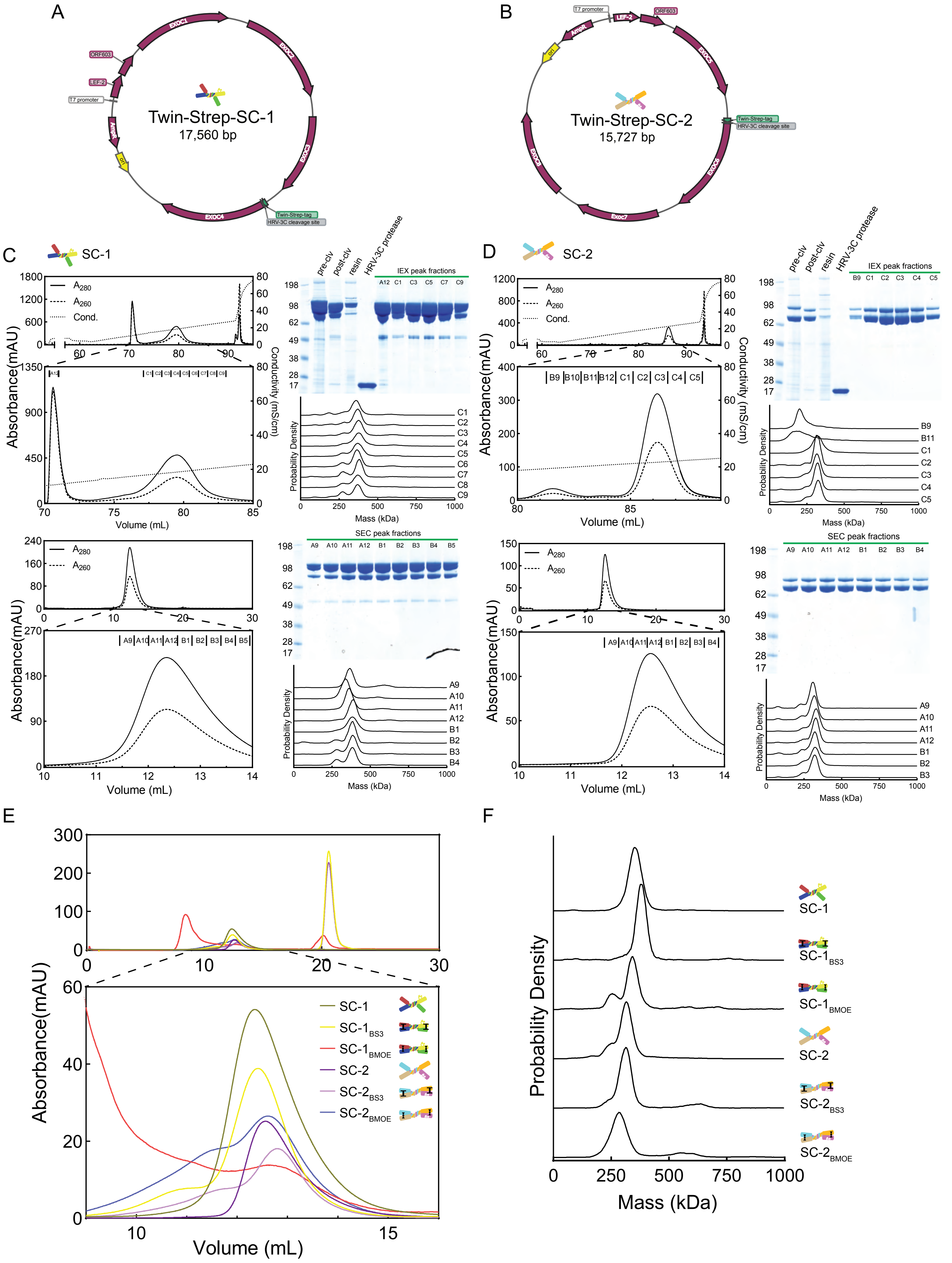

#### Figure S3.

A

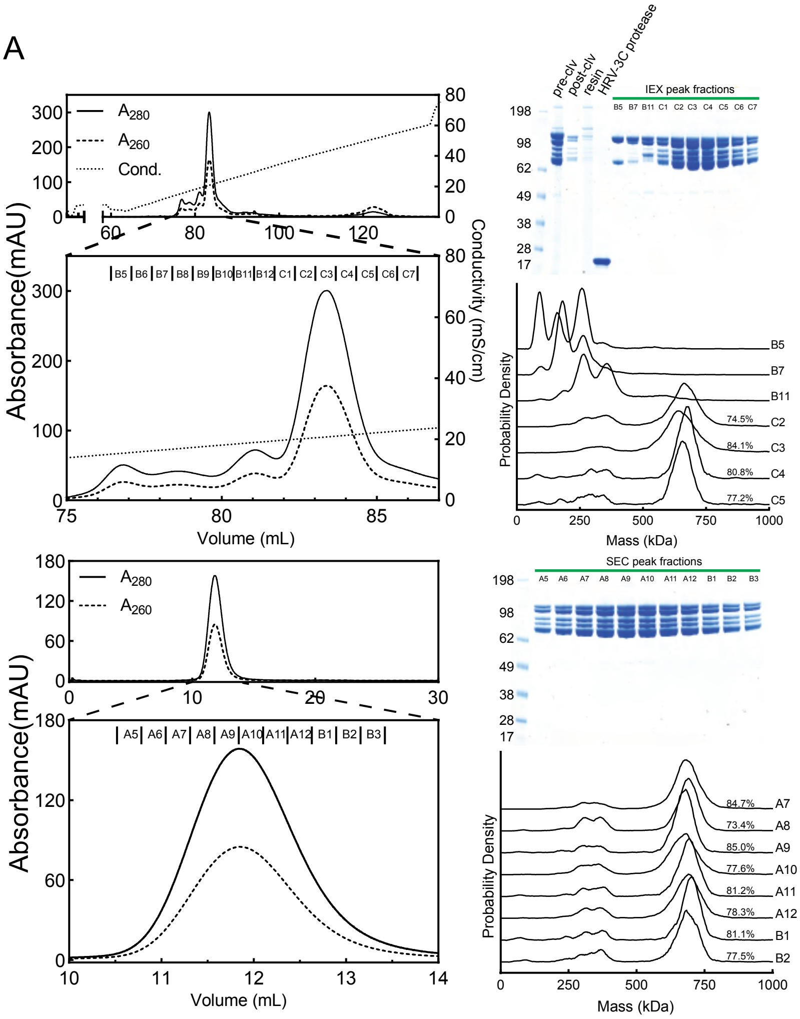


B

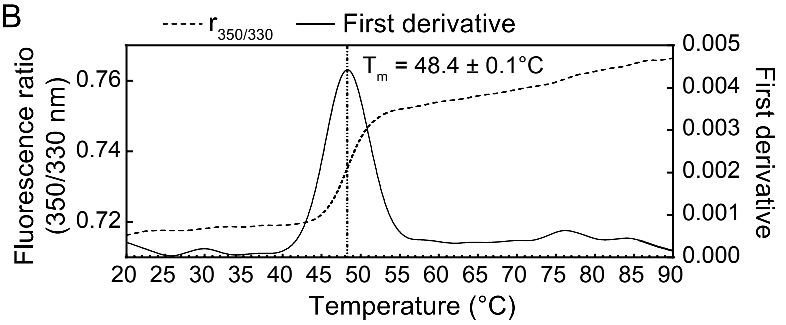


C

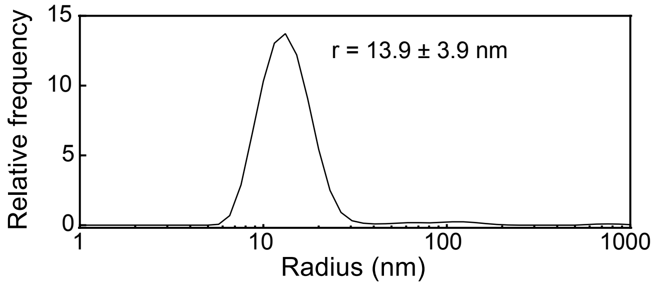

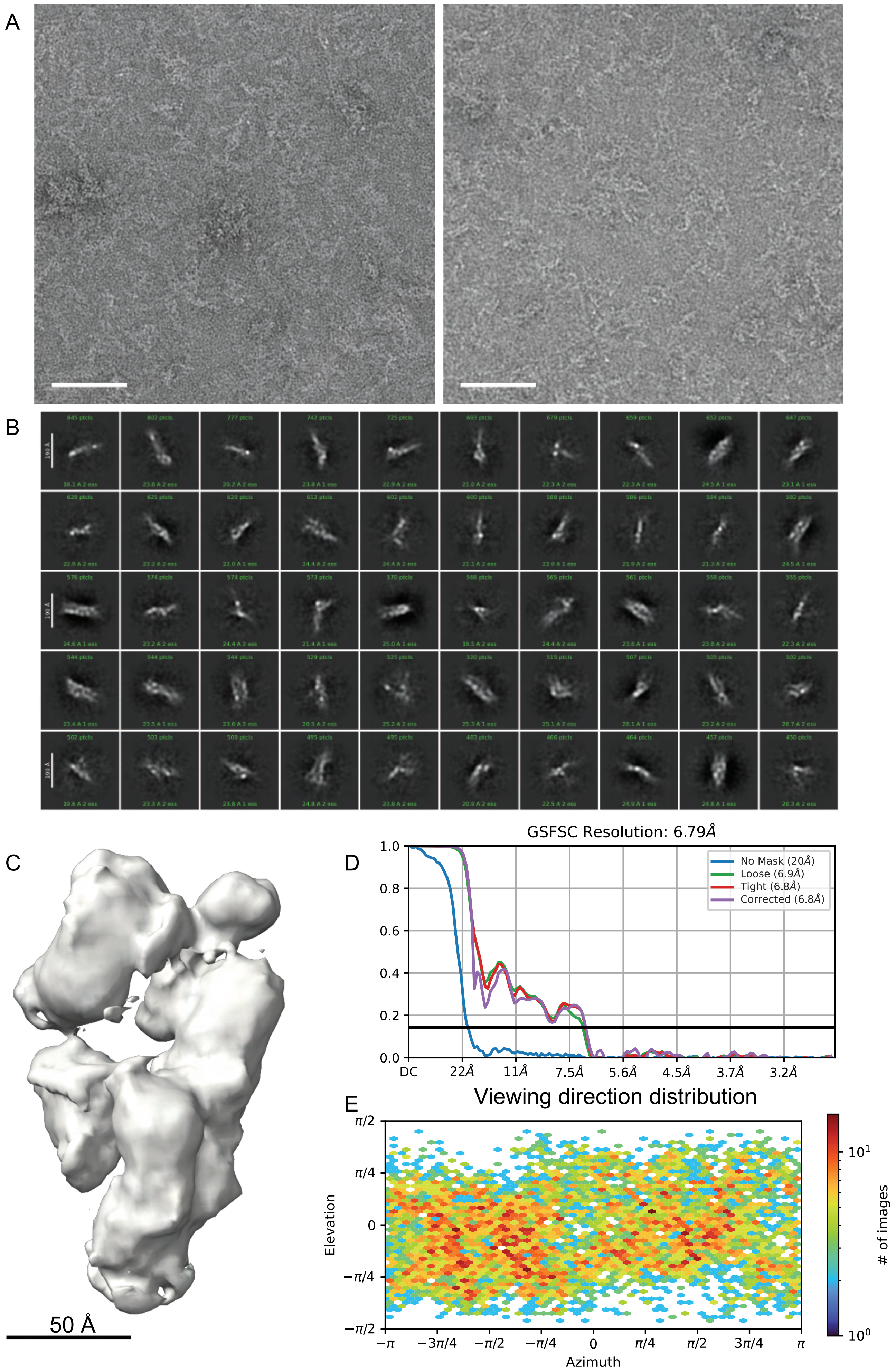

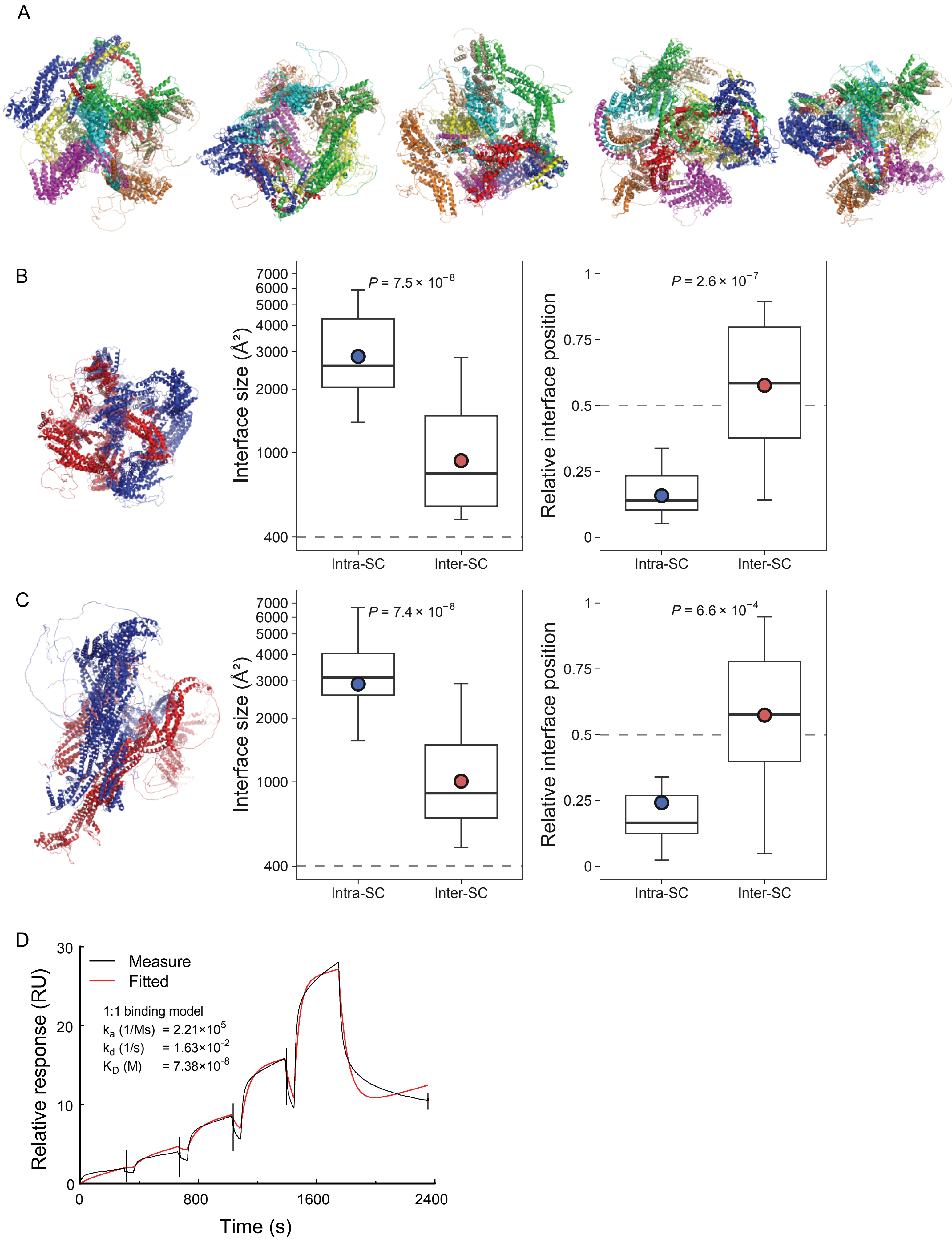

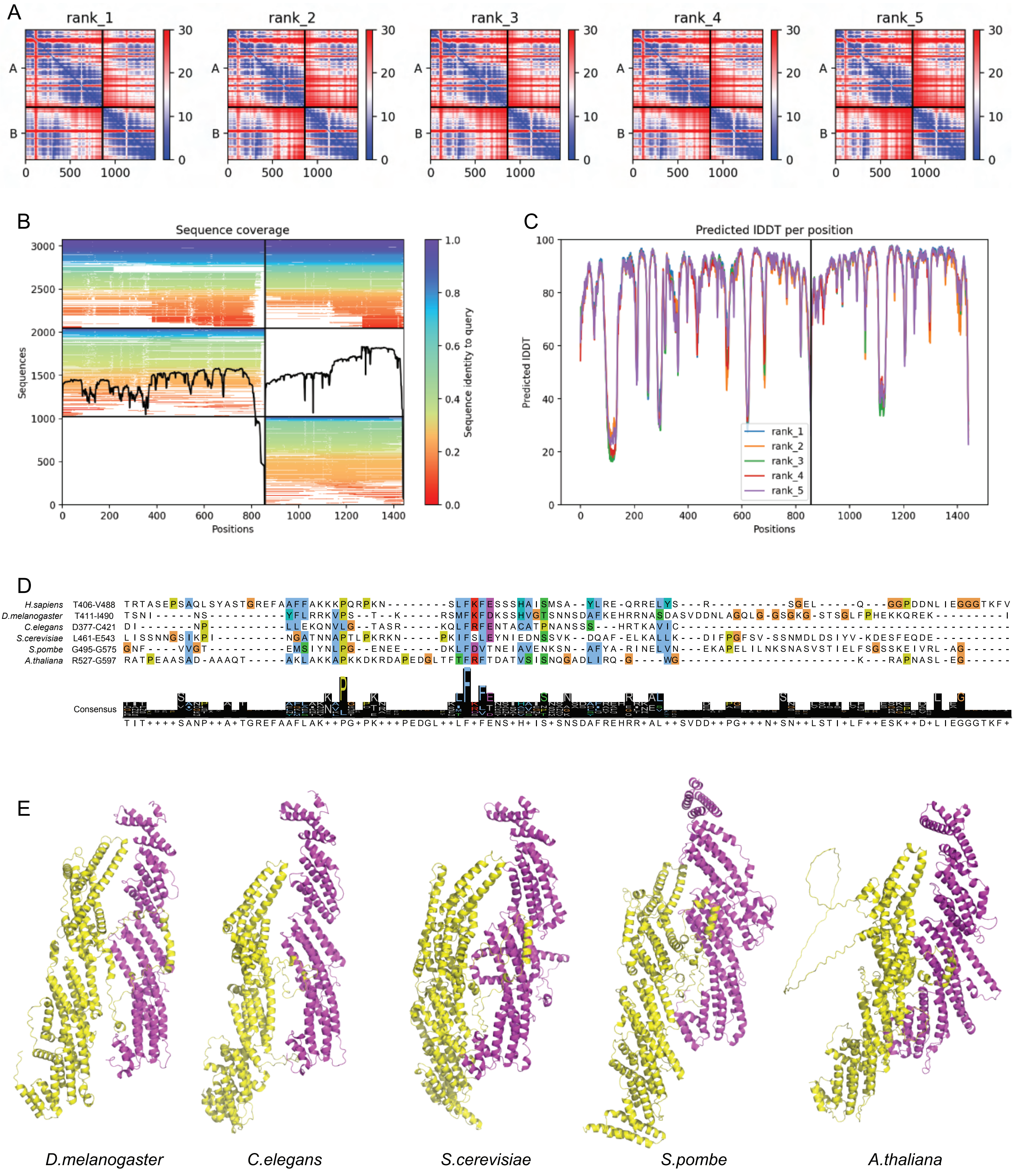

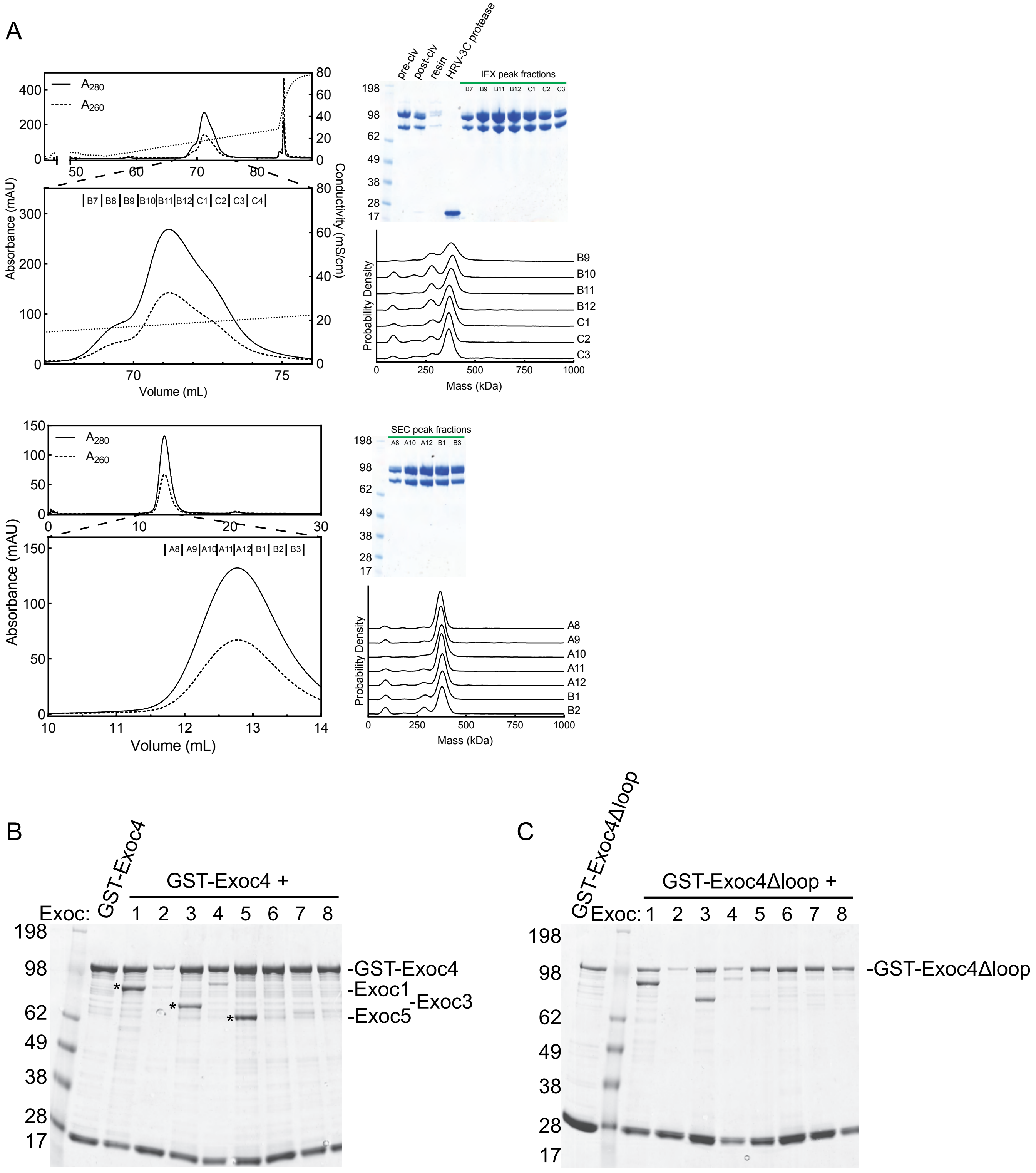
